# Coordinate Up-regulation of Both TK1 and TS Expression in Cycling Cells: Mechanistic Basis and Implications for Dual-Targeted Cancer Therapy

**DOI:** 10.64898/2026.01.02.697331

**Authors:** Wanxing Wang, Jiawei Zhang, Guangyou Duan, Yiyao Zhang, Guofeng Xu, Zhi Cheng, Shunmei Chen, Xue Yao, Xin Li, Shan Gao

## Abstract

Our previous study [2] has revealed that various types of cycling cells undergo rapid proliferation and division. Cycling cells can be broadly categorized into two groups: cycling immune cells (CICs) and cycling non-immune cells (CNICs). Cancer cells represent a special type of cycling cells. Based on a set of 26 marker genes, we proposed a simple method for the rapid identification of cycling-cell types. Comparisons across cycling-cell types uncovered novel gene-regulatory mechanisms underlying the cell cycle, development, and differentiation. Notably, we discovered that the coordinate up-regulation of both TS and TK1 expression is indispensable for securing a robust deoxythymidine triphosphate (dTTP) supply required for DNA replication in cycling cells. The compensatory interplay between salvage and *de novo* synthesis provides a plausible explanation for resistance to 5-fluorouracil (5-FU), a TS inhibitor for cancer chemotherapy. Consequently, blocking dTTP synthesis requires co-inhibition of TK1 and TS. For the first time, we present the crucial single-cell transcriptome evidence for this coordinate up-regulation, thereby establishing the mechanistic basis for developing TK1/TS as a dual target in cancer therapy.

## Introduction

Historically, rapid proliferation and division have been regarded as hallmarks of cancer cells. However, through the re-analysis of large-scale single-cell RNA sequencing (scRNA-seq) data [1], our previous study [2] has revealed that cycling cells also undergo rapid proliferation and division. Cancer cells represent a special type of cycling cells. Cycling cells can be broadly categorized into two groups: cycling immune cells (CICs), such as cycling myeloid cells (CMCs) [2], and cycling non-immune cells (CNICs), which mainly include stem cells and cancer cells. For instance, mesenchymal stem cells (MSCs), a type of CNIC, are implicated in the formation of cavernous hemangiomas, as reported in our previous study [3]. Our previous study has revealed that CICs rapidly expand in response to specific conditions, such as traumatic injury, pathogen infection, and cancer [2]. Given their dynamic expansion, CICs decisively influence both the course and outcome of the ensuing response. Therefore, accurate identification of cycling-cell types is the first step toward tracing their origins and deciphering their roles in both physiological homeostasis and pathological progression.

Our previous study [2] has established CICs as direct progenitors of immune cells and categorized them into two classes: CMCs and cycling lymphoid cells (CLCs). CMCs include a, d and m types, namely aCMC, dCMC, and mCMC, respectively, while CLCs include t and b types, namely tCLC and bCLC, respectively. These five types of CICs (aCMC, dCMC, mCMC, tCLC and bCLC) have been clearly defined in our previous study [2]. Additionally findings include: (1) aCMCs have potentials to differentiate or mature into microglia, monocytes, or macrophages, while dCMCs and mCMCs serve as progenitors for dendritic cells and mast cells, respectively; and (2) tCLCs are progenitors of T cells or natural killer cells, while bCLCs are progenitors of B cells. Our survey of published results from previous studies revealed that the accurate identification of different cycling-cell types remains challenging. They are often misidentified — most commonly as the mature cell types into which they differentiate.

Prior to the advent of scRNA-seq, accurate identification of cycling-cell types was infeasible. However, even with scRNA-seq, discriminating among CIC types remains neglected, particularly when their characteristics are not clearly defined. Fortunately, we developed a set of 68 marker genes, facilitating the identification of cycling cells. Then, we proposed a simple method for the rapid identification of each type of cycling cells. Based on large-scale scRNA-seq data mining, we systematically resolved cycling-cell types across tissues and conditions, and investigated their common features. In the present study, we demonstrate the utility of our method for the rapid and accurate identification of CICs and CNICs. We further report that that both TK1 and TS expression are coordinately up-regulated in cycling cells.

## Results

### Discriminating among CIC types in infected mouse livers

In our previous study [2]. we developed a set of 68 marker genes, facilitating the identification of cycling cells. Although these genes were initially identified in cancer tissue and conventionally regarded as oncogenes, they are highly expressed in all types of cycling cells and not specifically indicative of cancer. In subsequent studies, we refined the initial set of 68 genes to 26 genes. Based on these 26 genes, we proposed a simple method for the rapid identification of cycling cells, which in turn facilitates the identification of every remaining cell population in the dataset. Our method works in two steps: first, it identifies cycling cells using the 26 marker genes (**Figure 1A**); and second, it identifies each type of CICs based on two criteria: (1) it expresses marker genes of its corresponding mature immune cell type at a high level, typically slightly lower than that of the mature immune cell type (**Figure 1B**); and (2) its group is more closely located to its corresponding mature immune cell type, compared to other groups in the UMAP plot (**Figure 1C**). The second step can also identify each type of CNICs in a similar manner.

**Figure 1.**
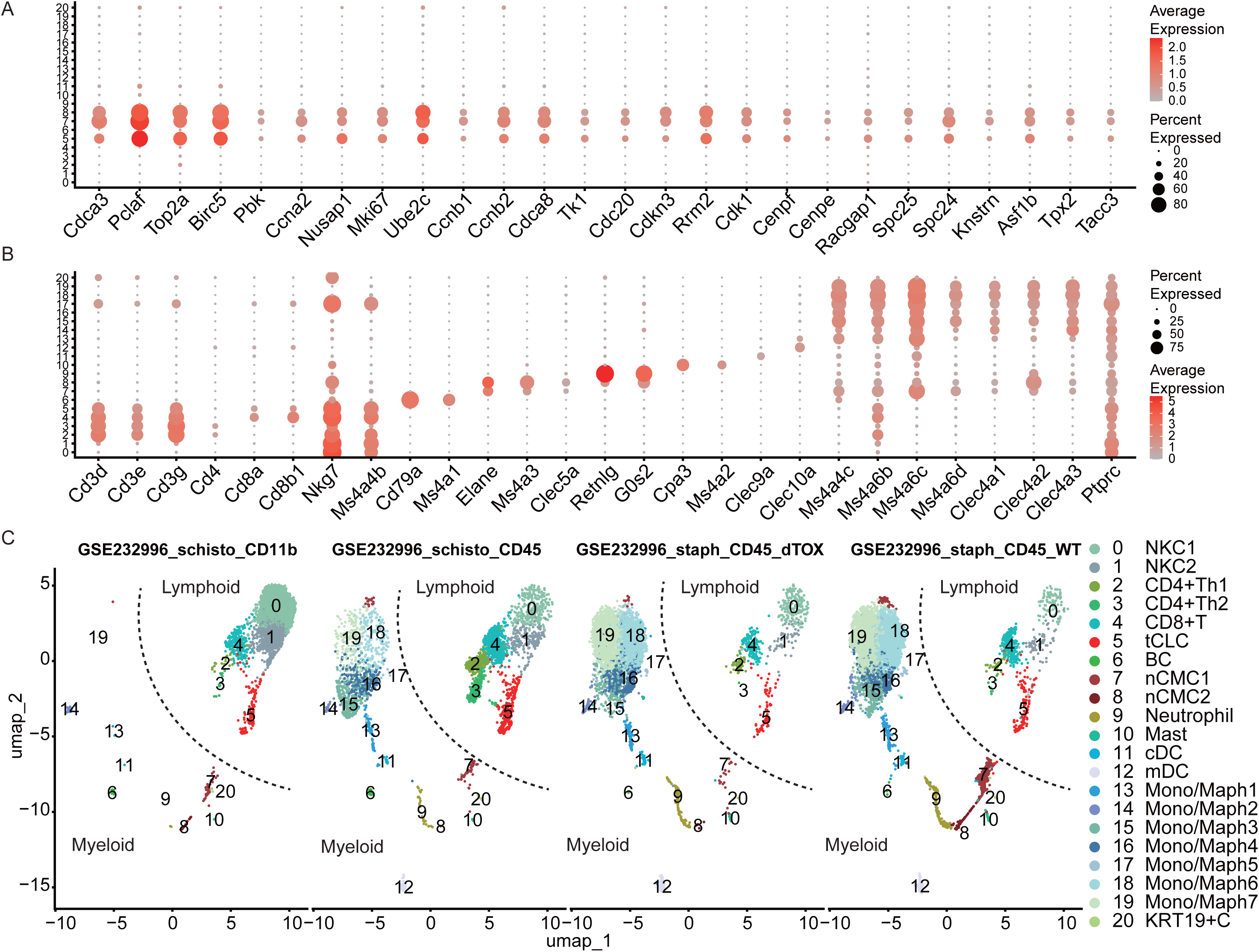
Discriminating among cycling-immune-cell types. From the scRNA-seq dataset (GEO: GSE232996), a total of 15,659 cells were clustered into 21 major groups, labeled from 0 to 20. NKC: natural killer cell; CD4+Th1: CD4 positive T helper cell type 1; CD4+Th2: CD8+T: CD8 positive T cells; tCLC: cycling lymphoid cell type t; nCMC: cycling myeloid cell type n; mDC: monocyte derived dendritic cell; cDC: CLEC9A positive dendritic cell; Mono/Maph: monocytes and macrophages; Mast: mast cell; BC: B cell; KRT19+C: KRT19 positive cell. **A**. 26 marker genes used to detect three groups of cycling cells, labeled 5, 7 and 8; **B**. 26 marker genes used to identify immune cells; **C**. The Uniform Manifold Approximation and Projection (UMAP) method was used to show 21 major groups under four conditions, including: (1) schisto_CD11B representing CD11B-captured mouse live cells infected by the parasite *Schistosoma mansoni*; (2) schisto_CD45 representing CD45-captured mouse live cells infected by the parasite *S. mansoni*; (3) staph_CD45_dTOX representing CD45-captured mouse live cells infected by the wild-type bacterium *Staphylococcus aureus*; and (4) staph_CD45_WT representing CD45-captured mouse live cells infected by the leukocidin-deficient *S. aureus*. A boundary (indicated by dash lines) was established between the nCMC1 and tCLC groups, segregating 20 of the 21 major groups into lymphoid and myeloid regions. Experiments under each conditions was repeated four times.

In the following paragraphs, we demonstrate the utility of our method for discriminating among CIC types by re-analyzing an infected mouse liver dataset from a previous study [4] as an example. This re-analysis clustered 15,659 cells into 21 major groups (**Figure 1C**) with adjusted parameters (**Materials and Methods**). Among these, three groups of cycling cells were identified based on expression profiles of the 26 marker genes (**Figure 1A**). Further analysis discriminated tCLCs from two groups of CMCs by three marker genes, *CD3D*, *CD3E* and *CD3G*. The two groups of CMCs, designated as nCMC1 and nCMC2, were further identified as progenitors of neutrophils, based on two criteria: (1) they highly expressed the neutrophil-specific marker gene *ELANE*, as well as *MS4A3*, a marker for early myeloid differentiation (**Figure 1B**); and (2) they were more closely located to the neutrophil group, compared to other groups in the UMAP plot (**Figure 1C**). These groups represent a novel type of CMCs, designated as the n type. Specifically, nCMC2 exhibited a closer relationship to neutrophils, as evidenced by the secondly highest expression of *RETNLG* and *G0S2*, while nCMC1 appeared to have a higher rate of rapid proliferation and division, with lower levels of *MS4A3*, compared to nCMC2.

To assist in identifying the remaining cell populations, a boundary was established between nCMCs and tCLCs, segregating 20 of the 21 major groups into two distinct regions (**Figure 1C**). The lymphoid region included tCLC and five other lymphoid groups, while the myeloid region included nCMC1, nCMC2, and 12 other myeloid groups. As an exception, the B cell (BC) group was mispositioned in the myeloid region due to its small size. The five lymphoid groups identified were the CD8 positive T cell (CD8+TC), two subtypes of CD4 positive T helper cells (CD4+Th1 and CD4+Th2), and two natural killer cell groups (NKC1 and NKC2). They were arranged in order of increasing distance from the tCLC group. In the present study, we defined a CD4/8 cell-type index (**Materials and Methods**) as the ratio between the average expression levels of *CD4* and *CD8A* within a group. Using this index, we confirmed the identities of CD8+TC, CD4+Th1, and CD4+Th2 based on their respective CD4/8 cell-type indices of -5.55, 3.85, and 4.87 (**Materials and Methods**). Among the 12 myeloid groups, the one closest to nCMC2 in the UMAP plot was the neutrophil group, while the second closest group was identified as mast cells by the marker gene *CPA3*. Of the remaining 10 myeloid groups, two were identified and named as the monocyte derived dendritic cell (mDC) and CLEC9A positive dendritic cell (cDC) groups, which were dispersed in distribution, while the other seven were monocyte or macrophage (Mono/Maph) groups, which were more densely packed and arranged in order of increasing distance from the nCMC2 group. The final group, characterized by high expression of ribosomal protein-encoding genes, was designated as KRT19+C due to its highly expressed gene *KRT19*.

This previous study [4] reported the expansion of *BIRC5* positive cells in mouse livers infected with wild-type bacterium *Staphylococcus aureus*, compared to those infected with leukocidin-deficient *S. aureus*. Unfortunately, the researchers were unable to discriminate among different types of cycling immune cells involved in the infection and immune responses. By re-analyzing their dataset, we correctly identified nCMCs and tCLCs (**Figure 1C**), both of which were included in the previously identified *BIRC5* positive cells. Subsequently, we determined that these *BIRC5* positive cells induced by *S. aureus* leukocidin are nCMCs, rather than tCLCs. Furthermore, we found that infection with the parasite *Schistosoma mansoni* led to a rapid expansion of tCLCs, rather than nCMCs. Although the previous study [4] reported that deletion of these *BIRC5* positive cells improves survival with *S. aureus* infection, it remains unclear whether nCMCs, tCLCs, or both play a major role in the immune response. Our re-analysis suggested that nCMCs and neutrophils, rather than tCLCs, play a major role in the immune response to *S. aureus* leukocidin infection during the 7-day post-infection period.

### Discriminating among closely related CIC types in tumor tissues

In the following paragraphs, we demonstrate the utility of our method for discriminating among closely related CIC types in tumor tissues by re-analyzing a breast cancer dataset from a previous study [5] as an example. This re-analysis clustered 49,973 cells into 20 major groups with adjusted parameters (**Materials and Methods**). Among these, two major groups were detected as cycling cells based on expression profiles of the 26 marker genes (**Figure 2A**). The cells in one group, which expressed immune marker genes (**Figure 2B**), were identified as CLCs, whereas those in the other group, which lacked immune marker gene expression, were conclusively inferred to be cancer cells. This inference was further confirmed by the cancer group’s closer proximity to the epithelial cell groups, compared to that of the CLC group (**Figure 2C**). Subsequently, a clear boundary was established between the CLC and cancer groups, segregating the 20 major groups into the immune and non-immune regions. With specific marker genes (**Figure 2B**), the 18 other major groups were identified. Among the 18 groups, six immune groups were further analyzed. Four of these six groups included cells that express both T cell- and B cell-specific marker genes, such as *CD3D*, *CD3E*, *CD3G*, *CD79A*, and *CD79B*. To quantify the relative abundance of T and B cells within these groups, we defined a T/B cell-type index (**Materials and Methods**) as the ratio between the average expression levels of *CD3D* and *CD79A* within a group. Using this index, the four groups were temporarily named as follows: TC (T Cell), T/BC (T/B Cell Mixed), B/TC (B/T Cell Mixed), and BC (B Cell). The CLC group closest to these four T/B groups was temporarily named as t/bCLC. Finally, the plasma group was identified based on the expression profiles of plasma cell-specific marker genes, such as *IGKC*, *IGLC2*, *IGHG1*, *IGHG3*, and *IGHG4*.

**Figure 2.**
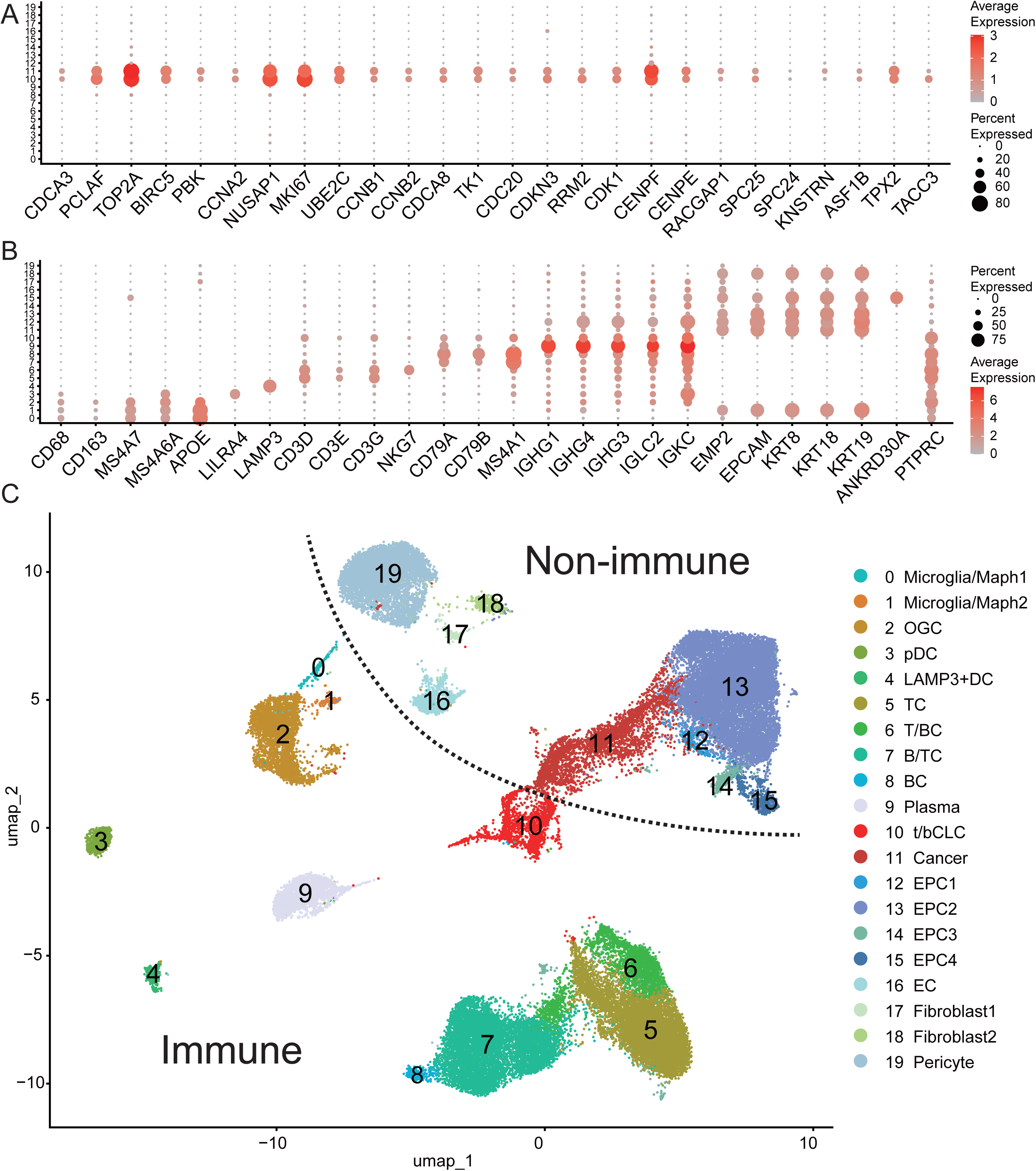
Discriminating among closely related CIC types. From the scRNA-seq dataset (GEO: GSE180286), a total of 49,973 cells were clustered into 20 major groups, labeled from 0 to 19. Microglia/Maph: Microglia or macrophages; OGC: Osteoclast-like giant cells; pDC: plasmacytoid dendritic cell; LAMP3+DC: LAMP3 positive dendritic cell; TC:T cell; T/BC:T or B Cell; B/TC: B or T Cell; BC: B cell; t/bCLC: cycling lymphoid cell type t or b; EPC: epithelial cell. EC: endothelial cell. **A**. 26 marker genes were used to detect two groups of cycling cells, labeled 10 and 11; **B**. 26 marker genes were used to identify types of major groups; **C**. A boundary (indicated by dash lines), named Nankai boundary [8], was established between the cancer and t/bCLC groups, segregating the 20 major groups into non-immune and immune regions.

A total of 21,625 cells from the six immune groups (TC, T/BC, B/TC, BC, t/bCLC and plasma) were selected for further analysis. These six groups were re-clustered into seven new groups, including cancer-cell, plasma, B-cell, T-cell, bCLC and tCLC groups and a mix group (**Supplementary file 1)**. Notably, the T-cell group was predominantly composed of double-negative T (DNT) cells, which are also referred to as CD3+CD4-CD8-T Cells [6]. Out of the 21,625 cells, 13,247 were CD3+ and 83.18% (11,019/13,247) of these were DNT cells—far exceeding the 55.74% (1,675/3,005) observed among CD3+ cells across six groups (NKC1, NKC2, CD4+Th1, CD4+Th2, CD8+T, and tCLC) in the mouse liver dataset (**Figure 1**). We further characterized the 11,019 DNT cells by their gene expression signature, which included the absence of *CD4*, *CD8A*, *CD8B*, *NKG7* and the high expression of *CD3D*, *CD3E*, *CD3G*, *GZMA*, *GZMK*, *GZMM*, and *IL7R*. Accordingly, the only tCLC group among the seven new groups was conclusively inferred to be the precursor of these DNT cells by their shared marker genes and the relative distance in the UMAP plot (**Supplementary file 1)**. Consequently, this tCLC group was designated as the DNtCLC group. Among three myeloid groups related to monocytes or macrophages (**Figure 2C**), one was predominantly composed of osteoclast-like giant cells (OGCs), which were characterized by their high expression of *CD68*, *CD86*, and *CD163* [7], compared to the other two groups. However, osteoclast-like giant type cycling myeloid cells (oCMCs) was not detected using this dataset, but was identified in seven other breast cancer datasets (**Supplementary file 1)**. Additionally, tCLCs, oCMCs, bCLCs, and aCMCs were detected across multiple types of breast cancer using these datasets, showing no specificity to particular cancer types.

### The differentiation of nCMCs and tCLCs

Our previous study [2] revealed that aCMCs primarily differentiate or mature into microglia in the CNS and may also differentiate into monocytes or macrophages in other tissues; by extension, every CMC or CLC type is poised to differentiate into the corresponding mature immune cell of its own lineage. Figure 1 shows that nCMCs differentiate into neutrophils, as neutrophil is the sole mature population adjacent to the nCMC1 and nCMC2 groups. In contrast to nCMCs, tCLCs sit at a lineage crossroads, as their (*CD8A*, *CD8B*, *NKG7* and *MS4A4B*) are shared by CD8+T and NK cells, while the absence of CD4+ excludes a CD4+TC fate. To resolve the differentiation routes of nCMCs and tCLCs, we performed pseudotime analysis (**Materials and methods**) on two subsets of the mouse liver dataset (GEO: GSE232996). Subset 1 contained 946 cells from the nCMC1, nCMC2 and neutrophil groups, whereas subset 2 contained 2,698 cells from the tCLC, CD8+TC, CD4+Th1, and CD4+Th2 groups (**Figure 2C**). Using the top 497 highly variable genes (HVGs), we reconstructed a linear differentiation path in subset 1 (**Supplementary file 1**) that originates in nCMC1 (branch a, early pseudotime), transits through nCMC2 (intermediate pseudotime) and terminates in mature neutrophils (branch b, late pseudotime). The uniform distribution of nCMC2 cells across both branches confirms their transitional identity, forming a continuous bridge between the progenitor and terminally differentiated neutrophils. Using the top 2,000 HVGs, pseudotime analysis of subset 2 (**Supplementary file 1**) placed tCLCs at the root of a trajectory that terminates in CD8+TCs, with CD4+Th1 and CD4+Th2 cells branching off separately—indicating a CD8+TC fate rather than an CD4+TC fate for tCLCs.

### Discovery of up-regulated expression of both TK1 and TS

By a large-scale mining of scRNA-seq data in the DISCO database [8], we confimed that the 26 marker genes are expressed in every cycling-cell type we identified, constituting the most universal feature of cycling cells (**Supplementary file 1**). Among them, *TK1* and *RRM2* encode enzymes in deoxyribonucleotide (dNTP)-synthesis pathways. Notably, TK1 functions in the salvage pathway for deoxythymidine triphosphate (dTTP) synthesis, which has received far less attention than *de novo* dNTP-synthesis pathways in which *RRM2* functions (**Figure 3A**). These *de novo* pathways generate dNTPs by reducing ribonucleotides (NTPs) to their deoxy forms. All dNTP-synthesis enzymes have been mapped to their encoding genes including *RRM1*, *RRM2*, *RRM2B*, *DCK*, *DGUOK*, *CTPS1*, *CTPS2*, *DCTD*, *CDADC1*, *DUT*, *TYMS*, *TK1* and *TK2.* Among them, *RRM1*, *RRM2* and *RRM2B* encode ribonucleotide reductase (RNR) [9] and *TYMS* encodes thymidylate synthase (TS) [10]. The functions of these enzymes have been systematically characterized. However, their regulatory dynamics were unclear. Comparison across every cycling-cell type we identified led us to discover that the expression of *RRM1*, *RRM2*, *DCK*, *DUT*, *TK1* and *TYMS* are coordinately up-regulated (**Figure 3BC**). For example, in aCMCs [2], their relative abundance are 5.79 (*RRM1*), 18.65 (*RRM2*), 5.67 (*DCK*), 3.91 (*DUT*), 1.37 (*TK1*) and 1 (*TYMS*), representing 1.55-, 2.58-, 1.40-, 1.61-, 2.35- and 1.81-fold increases over non-cycling cells, respectively (**Supplementary file 1**). Coordinated up-regulation of these enzymes enables rapid expansion of the dNTP pool within a narrow time window to meet the transient surge in dNTP demand for DNA replication, playing an indispensable role in cell-cycle progression.

**Figure 3.**
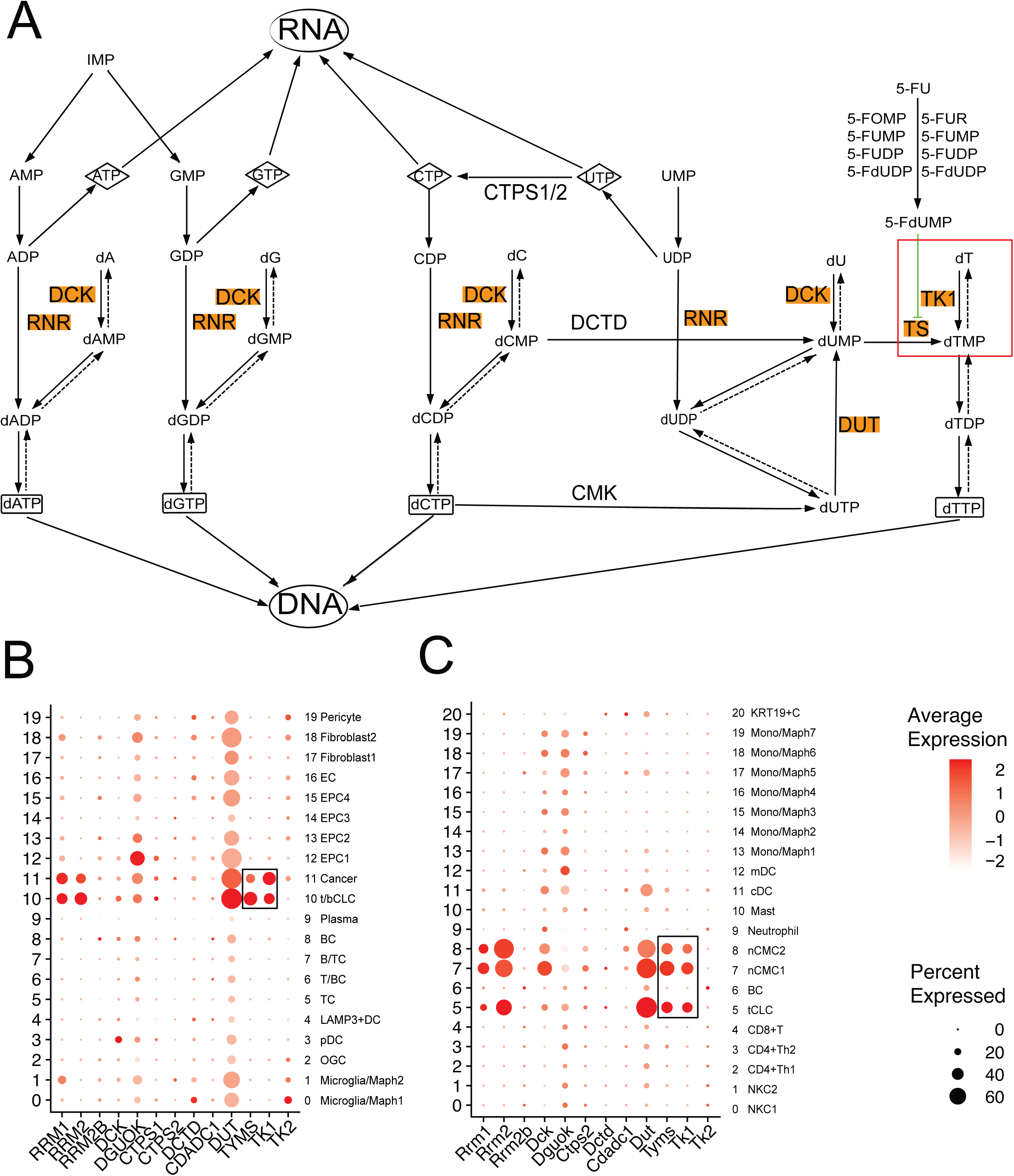
TK1 and TS in dNTP-synthesis pathways. **A**. The salvage pathway of dTTP synthesis has been integrated into the NTP/dNTP conversion map and the up-regulated core enzymes include RNR, DCK, DUT, TK1 and TS (in orange color); **B**. The expression of human *RRM1*, *RRM2*, *DCK*, *DUT*, *TK1* and *TYMS* are up-regulated in two groups of cycling cells (t/bCLC and cancer), out of 20 major groups clustered using a total of 49,973 cells from the scRNA-seq dataset (GEO: GSE180286); **C**. The expression of mouse *Rrm1*, *Rrm2*, *Dck*, *Dut*, *Tk1* and *Tyms* are up-regulated in three groups of cycling cells (tCLC, nCMC1 and nCMC2), out of 21 major groups clustered using a total of 15,659 cells from the scRNA-seq dataset (GEO: GSE232996). 5-FU: 5-Fluorouracil; CTPS1/2: CTP synthase 1/2; CMK (*CDADC1*): cytidine/uridine monophosphate kinase 1; RNR: ribonucleotide reductase; DCTD: deoxycytidylate deaminase; DCK: deoxycytidine kinase; DUT: dUTP diphosphatase; TK1/2: thymidine kinase 1/2; TS (*TYMS*): thymidylate synthase; DGUOK: deoxyguanosine kinase. RNR is encoded by *RRM1*, *RRM2* and *RRM2B*. TK2 and DGUOK are located in mitochondrion.

Analysis of the dNTP-synthesis pathways revealed that coordinate up-regulation of both TK1 and TS expression is indispensable for securing a robust dTTP supply (**Figure 3A**): TS plays a critical role in the *de novo* synthesis by reductively methylating deoxyuridine monophosphates (dUMPs) to deoxythymidine monophosphates (dTMPs), whereas TK1 launches the salvage synthesis by phosphorylating deoxythymidine (dT) to dTMP. Consequently, up-regulated expression of either enzyme compensates for the dTTP shortfall imposed by inhibition of the other, suggesting a potential resistance mechanism of 5-fluorouracil (5-FU), a TS inhibitor for cancer chemotherapy. Although 5-FU has been widely used since the 1950s to treat a broad spectrum of solid tumors including colorectal, breast, and lung cancers, resistance to this drug remains a formidable challenge. Although many mechanisms have been proposed [11], only a narrative review has begun to implicate dTTP salvage synthesis as a contributing factor [12]. However, the relative contributions of salvage and *de novo* dTTP synthesis to the transient surge in dNTP demand was unknown. At first glance—judging only by the expression levels and fold-changes of *TK1* and *TYMS* (**Supplementary file 1**)—the two pathways appear to contribute equally in cycling cells. Yet, *de novo* synthesis is limited by numerous additional enzymes — beyond the up-regulated RNR, DCK, DUT and TS — whose expression are not significantly up-regulated in cycling cells (**Figure 3**). In contrast, salvage synthesis relys solely on TK1. Consequently, salvage synthesis is markedly more efficient than *de novo* synthesis. If we further consider that TK1 is limited by the dT pool, which continuously declines as DNA replication proceeds, the salvage pathway along can not maintain prolonged dTTP supply. Collectively, salvage synthesis primarily delivers the rapid, initial burst of dTTPs, whereas *de novo* synthesis sustains the longer-term supply during DNA replication. During 5-FU therapy, dying or dead cells release dTs for salvage synthesis, allowing cancer cells to withstand or escape *de novo* inhibition. Consequently, blocking dTTP synthesis requires co-inhibition of TK1 and TS.

Among the 26 marker genes, *TK1* is the only one encoding a membrane protein, and its structure switches from a dimer in bacteria, plants and *Dictyostelium* to a tetramer in animals [13]. Currently, TK1’ up-regulated expression in cancer cells or tissues is attracting growing interest, and it has been proposed as a therapeutic target [14]. Our scRNA-seq data analysis now provided the missing crucial evidence at the gene-expression level: the coordinated up-regulation of multiple enzymes involved in dNTP-synthesis established the mechanistic basis for developing TK1/TS as a dual target in cancer therapy. To develop TK1 inhibitors, two critical issues must be confronted: (1) up-regulated expression of TS could compensates for the dTTP shortfall imposed by inhibition of the TK1, which can be circumvented by co-inhibition of both TK1 and TS; and (2) any therapeutic strategy targeting TK1 on cancer cells will inevitably hit CICs as well, yet up-regulated expression of TK1 in CICs remained unnoticed in previous studies. Earlier work has shown that TK1 loses its cell cycle control in cancer cells, and a fraction of TK1 can translocate to the plasma membrane, forming the so-called membrane-bound TK1 (mTK1) [14]. As both cancer cells and CICs are cycling cells, mTK1 should be present on both, enabling antibody-based detection in our following experiments. Using the MC38 tumor-bearing mouse model (**Materials and Methods**), we demonstrated that the fraction of CD8+T cells displaying mTK1 rose from 0.471 % in controls to 2.64 % in spleens of tumor-bearing mice (**Figure 5**), validating our prediction. These results confirmed that targeting mTK1 hits both cancer cells and CICs that display up-regulated expression of TK1. Fortunately, CICs constitute only a minute fraction of the immune cells, leaving the vastly larger pool of non-cycling immune cells intact to sustain anti-tumor immunity. On the other hand, given that almost all these CICs are tumor-elicited, they probably play tumor-promoting roles. Therefore, TK1/TS constitute a dual target whose co-inhibition simultaneously suppresses tumor growth and tumor-associated inflammation. Moreover, our confirmation of mTK1 on the cell surface provides a rationale for antibody-based targeting in future drug design.

## Conclusion and Discussion

The present study demonstrated the utility of our method for rapid and accurate identification of CICs and CNICs. However, due to the limitations of current scRNA-seq technologies, many important marker genes (*e.g.*, *CD4*, *CD8A* or *CD8B*) are detected at exceedingly low expression levels, posing challenges to the accurate identification of CIC subtypes such as DNTtCIC. Moreover, the origins and exact roles of most CIC types in various physiological and pathological processes remain largely unknown. Our previous study [2] revealed that CICs undergo rapid expansion in response to various stimuli, such as traumatic injury, pathogen infection, and cancer, mirroring the behavior of their corresponding immune cell types. This finding raises two scientific questions: 1), whether the expansion of CICs arises primarily from proliferation of a pre-existing and self-sustained pool or transition of their corresponding mature immune cells; and 2), whether the subsequent expansion of immune populations arises primarily from proliferation of resident immune cells or differentiation of their CIC progeny. Beyond these two scientific questions, comparisons across cycling-cell types will help researchers discover novel gene-regulatory mechanisms underlying the cell cycle, development, and differentiation.

To date, no inhibitors targeting human or mammalian TK1 have been approved for marketing or reported to be in development. Previous studies and current clinical strategies have targeted TS alone, primarily because the coordinate up-regulation of both TK1 and TS expression remained unrecognized. Re-examination of the integrated map (**Figure 3A**) confirmed that no other dTTP-synthesis pathway exists beyond the *de novo* and salvage arms. Therefore, their co-inhibition can completely block dTTP synthesis. Our experiments suggested that co-inhibition of TK1/TS simultaneously suppresses tumor growth and tumor-associated inflammation. This benefit, however, raises a critical question—could the same anti-inflammatory activity compromise anti-tumor inflammation during therapy? Our experiments confirmed that CICs constitute only a minute fraction of the immune cells (**Figure 4**), leaving the vastly larger pool of non-cycling immune cells intact to sustain anti-tumor immunity. Beyond oncology, co-inhibition of TK1/TS may therefore offer a general strategy to suppress inflammation in immune-driven disorders.

**Figure 4.**
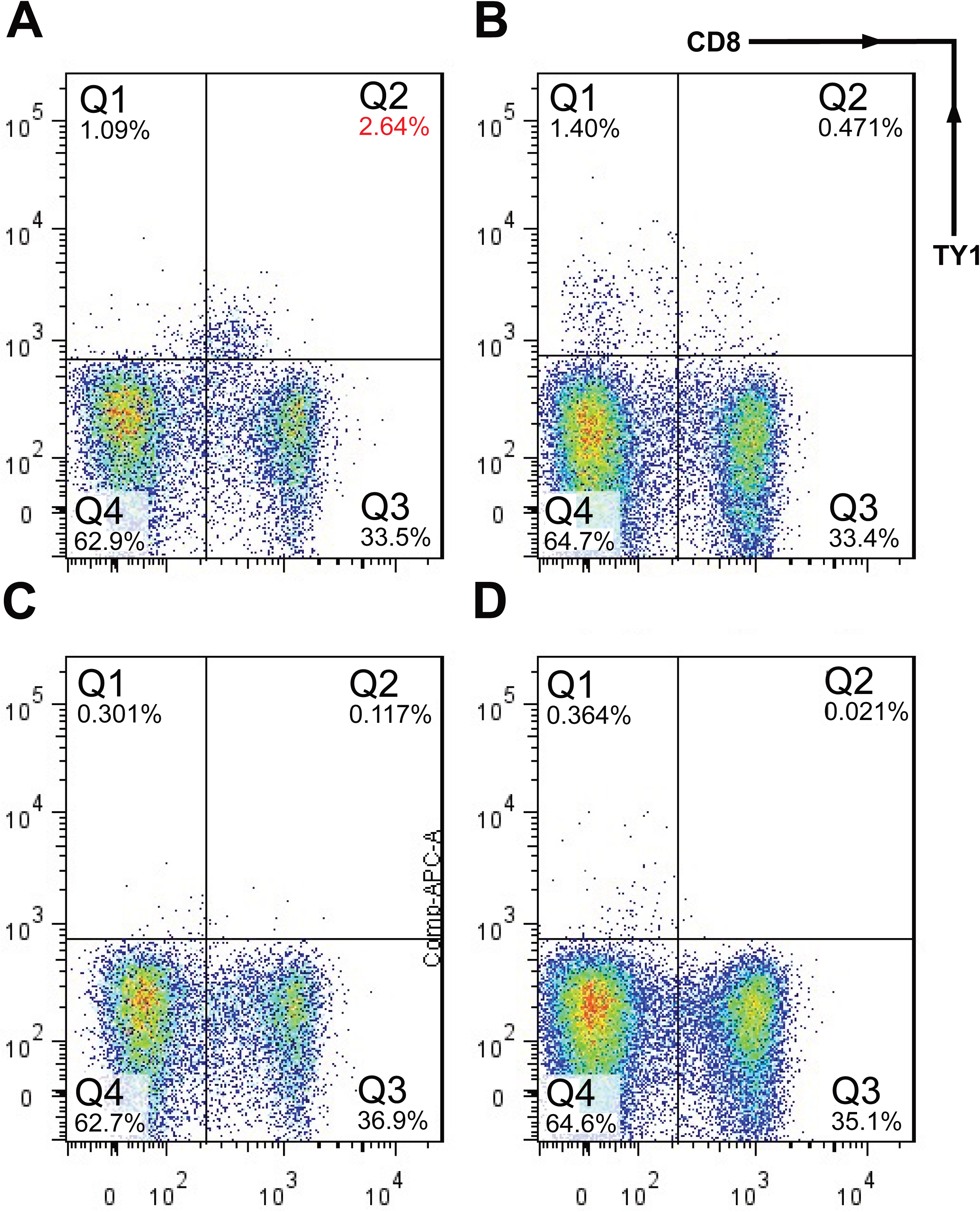
TK1 positive tCLCs from spleens of MC38 tumor-bearing mice. 110,359, 123,388, 109,946 and 128,846 splenocytes were sequentially gated as follows: FSC-W vs. FSC-H (singlets), SSC-W vs. SSC-H (singlets), FSC-A vs. Zombie+ (live cells), SSC-A vs. CD45+ (leukocytes), and CD3+ (T cells). The final CD45+CD3+ populations are displayed as CD8 (x-axis) versus TK1 (y-axis): A. 13,498 cells from spleens of tumor-bearing mice; B. 23,142 cells from controls; C. 13,637 cells from spleens of tumor-bearing mice; D. 28,605 cells from controls.

Rapid proliferation and division is the immutable hallmark of every cancer type, and the attendant surge in dNTP demand is their Achilles heel. Regardless of acquired drug resistance or immune-evasion tactics, rapid proliferation and division obliges cancer cells to expand their dNTP pools; consequently, no malignant cell can escape the dTTP depletion inflicted by co-inhibition of TK1/TS. Up-regulation of TK1 expression has been repeatedly observed in drug-resistant or immune-evading samples, indicating that these surviving cancer cells are still vulnerable to TK1 inhibition. For example, a recent phase-II trial reported that up-regulation of TK1 expression predicts resistance to immune checkpoint plus tyrosine kinase inhibition in renal cell carcinoma [15]. As up-regulation of TK1 expression is present in every cycling-cell type we identified, blocking dTTP synthesis could be developed into a universal, once-and-for-all solution to drug resistance and immune evasion across all cancer types.

## Materials and Methods

### All datasets and analysis

In the present study, the metadata of scRNA-seq data were downloaded from the DISCO database (https://www.immunesinglecell.org/) [8], which included information on 122,093,310 single cells of 243,608 cell types from 19,762 samples, as of March 8th, 2025. A mouse liver dataset (GEO: GSE232996) from a previous study [4] and a breast cancer dataset (GEO: GSE180286) [5] were used for demonstration. Each gene expression matrix was downloaded and re-analyzed by the same pipeline with different parameters, which was detailed described in our previous study [2]. The R package Seurat v4.1.0 and other R packages (*e.g.,* ggplot2) were used for scRNA-seq data analysis on R v4.1.3 [16]. To improve the data analysis, the TU (Total Ubiquitous) method in the R package NormExpression [17] was used to normalize gene expression data and the R package Harmony v4.1.0 [18] was used to remove the batch effects. The R package Monocle [19] v2.8.0was used to estimate pseudotemporal paths.

Cell-type identification started in two steps: cycling cells were first delineated by the 26-marker gene set (**Figure 1A**), and individual cycling-cell-type identities—including CICs, cancer cells, or stem cells—were then assigned using the same dual-criterion framework (**Results**). Non-cycling cells were subsequently identified by further analysis of its more top featured genes, or even all DE genes in its gene-expression signature using the Metascape website (https://metascape.org/gp) [20] a web-based portal designed to provide a comprehensive gene list annotation and analysis resource. All cell-type identifications were carried out only to the major groups and were not further subdivided into subgroups, except for two scRNA-seq datasets (GEO: GSE180286 and GSM5354533). CD4/8 index was designed to differentiate CD4 positive cells from CD8 positive cells. It is calculated as the base-2 logarithm of the ratio of *CD4* expression levels to *CD8A* expression levels.

### Cell and animal experiments

The experiments were conducted with the approval and consent of the Ethics Committee of Nankai University (No: 2022-SYDWLL-000079), which ensures that the research adheres to ethical guidelines and principles. The C57BL/6 mice used in the present study were purchased from Vital River Laboratory 110 Animal Technology Co., Ltd (Beijing, China, Permission Number: SCXK (Jing)-2016-0006). To construct the MC38 tumor-bearing mouse model, 1.25 x 10^6^ MC38 cells were injected into C57BL/6 mice (∼10-weeks, ∼20-25 g). Mice were euthanized when their tumor volumes reach 1500 mm³, as described in a previous study [21]. The isolation of splenocytes from each mouse was performed as follows: (1) swab the abdominal fur of the mouse with 70 % ethanol, and open the peritoneal cavity with sterile scissors on a clean dissection board; (2) lift the spleen with forceps after gently free it, and place it in a 15 mL tube containing 5 mL ice-cold sterile PBS; (3) transfer the spleen to a 70 μm cell strainer (pre-rinsed with 1 mL PBS) placed in a six-well plate, and then grind the tissue with the plunger of a sterile 1 mL syringe until only connective tissue remains; (4) rinse the strainer with 1 mL PBS to recover the residual cells; (5) suspend the cells collected from steps 3 and 4, centrifuge at 400 × g, 4 °C for 5 min, and discard the supernatant; (6) repeat step 5 to wash the cells; (7) resuspend the pellet in 5 mL ice-cold RBC lysis buffer, incubate on ice for 5 min, then terminate lysis by adding 10 mL sterile PBS; (8) Centrifuge the lysed cell suspension at 400 × g, 4 °C for 5 min, and discard the supernatant; (9) resuspend the cells in 5 mL PBS, and filter through a 40 μm cell strainer to remove clumps; and (10) Centrifuge the flow-through and collect the cells. Splenocytes were isolated from three tumor-bearing mice (tumor group) and three tumor-free controls (control group), and pooled to generate one tumor sample and one control sample. The cells were stained using Zombie NIR™ Fixable Viability Kit (BioLegend, China) and fluorochrome-conjugated antibodies, then counted on an LSRFortessa flow cytometer (BD, USA). FlowJo software was used for data analysis with the free functions of the software. Fluorochrome-conjugated antibodies (BioLegend, China) include: anti-CD45 (cat: 103108), anti-CD3 (cat: 100218), and anti-CD8 (cat: 100751). Particularly, anti-TK1 (cat: R21614) was purchased from Zen-Bioscience, China.

## Supplementary information

Supplementary file 1 (s1.docx)

## Declarations

### Ethics approval and consent to participate

Not applicable.

### Consent to publish

Not applicable.

### Availability of data and materials

All important data in the results is provided in the supplementary files.

### Authors’ contributions

Shan Gao conceived the project. Shan Gao and Xin Li supervised the present study. Jiawei Zhang, and Wanxing Wang collected the samples and conducted the experiments. Guangyou Duan and Yiyao Zhang performed programming. Guofeng Xu and Shunmei Chen analyzed the data. Zhi Cheng performed statistics and plotting. Shan Gao drafted the main manuscript text. Shan Gao and Xue Yao revised the manuscript.

## Acknowledgments

We are grateful for the help from Nanxin Gong from College of Life Sciences at Nankai University. This manuscript was online as a preprint on Dec. 10th, 2025 at.

