## Supplementary material for "Coordinate Up-regulation of Both TK1 and TS Expression in Cycling Cells: Mechanistic Basis and Implications for Dual-Targeted Cancer Therapy": s1

### Supplementary file 1

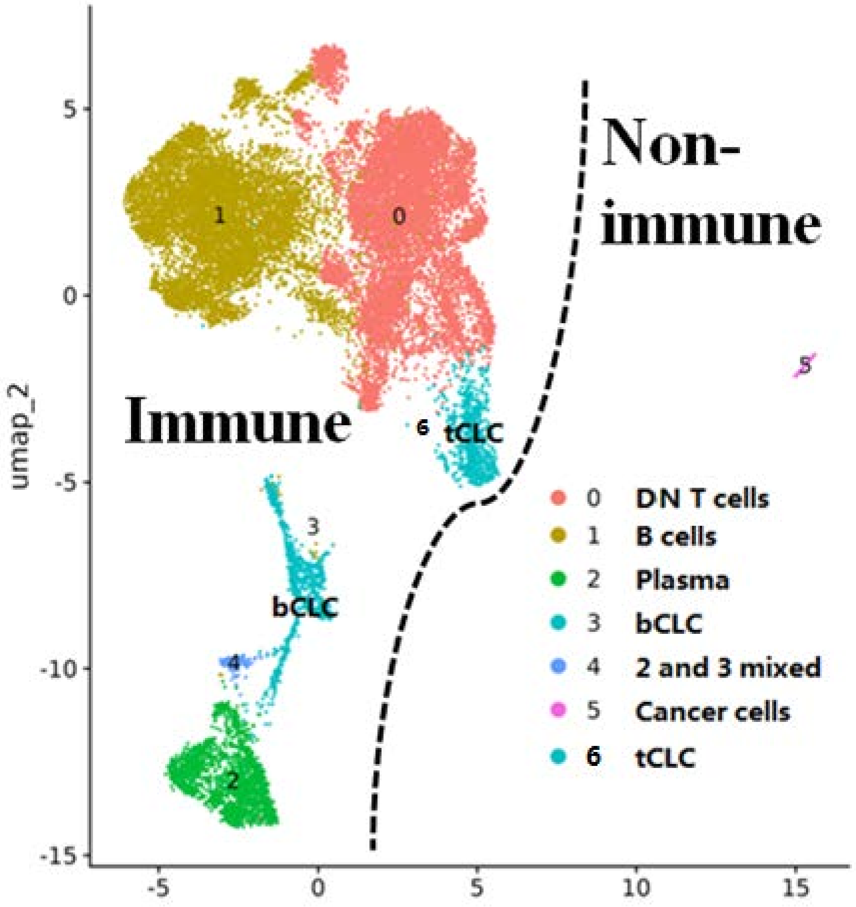


**Fig S1. Seven new groups from further clustering analysis**

A total of 21,625 cells from these six major groups (GEO: GSE180286) were selected for further clustering analysis. This analysis resulted in seven new groups, which were designated as groups 0 to 6. Notably, a group (in blue color), originally identified as the t/bCLC group, was divided into two separate groups: the tCLC group and the bCLC group. DNT cells: double-negative T cells; tCLCs: T subtype cycling lymphoid cells; bCLCs: b subtype CLCs.

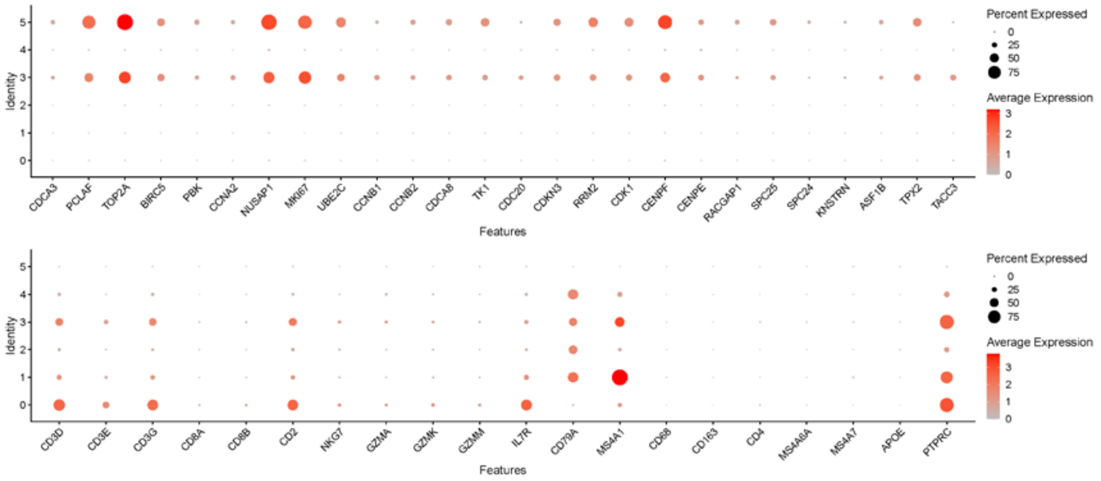


**Fig S2. Six new groups identified using cell-specific markers**

Six cell groups, numbered from 0 to 5, correspond to those in **Figure 1** on a one-to-one basis. Group 3 includes tCLCs and bCLCs. The tCLCs are predominantly composed of double-negative T subtype cycling lymphoid cells (DNtCLCs).

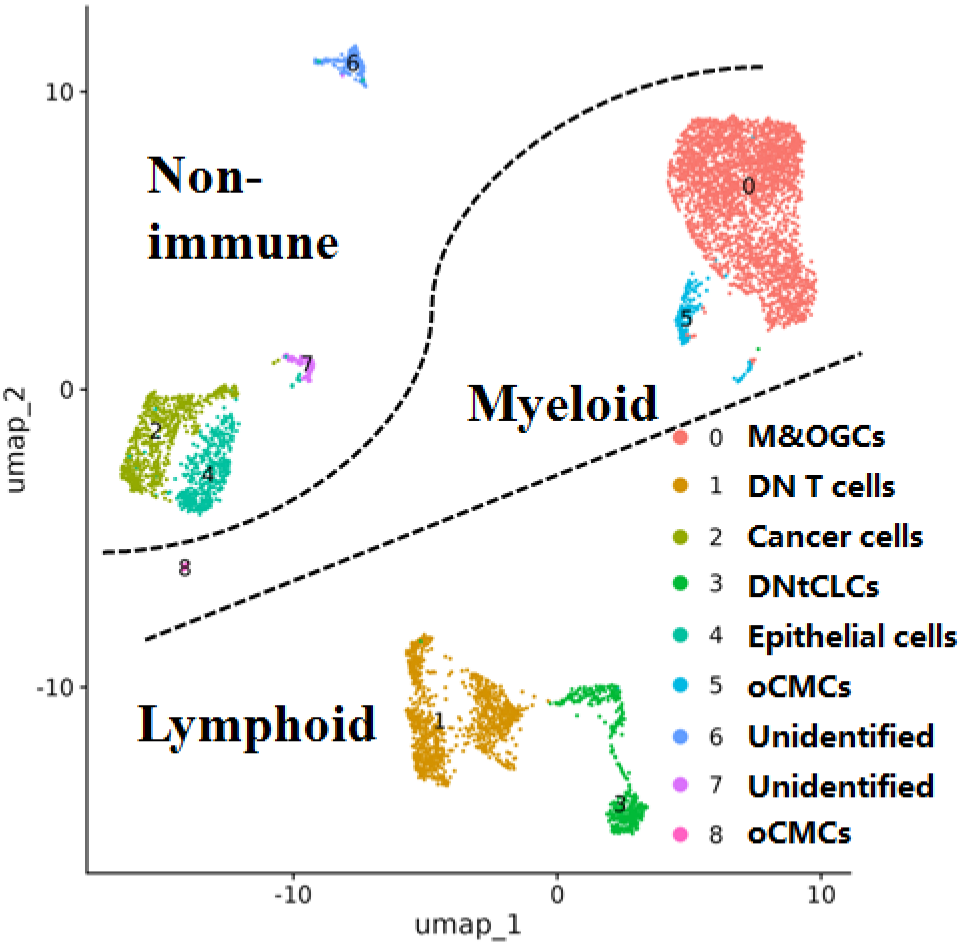


**Fig S3. Three types of cycling cells identified in a sample**

A total of 5,619 cells in a tumor tissue (GEO: GSM5354533) were clustered into nine major groups, which were designated as groups 0 to 8. Three types of cycling cells (Cancer cells, DNtCLCs, and oCMCs) were identified in this sample. M&OGCs: Monocytes or macrophages that are predominantly composed of osteoclast-like giant cells; oCMCs: osteoclast-like giant subtype cycling myeloid cells; DNT cells: double-negative T cells; DNtCLCs: double-negative T subtype cycling lymphoid cells.

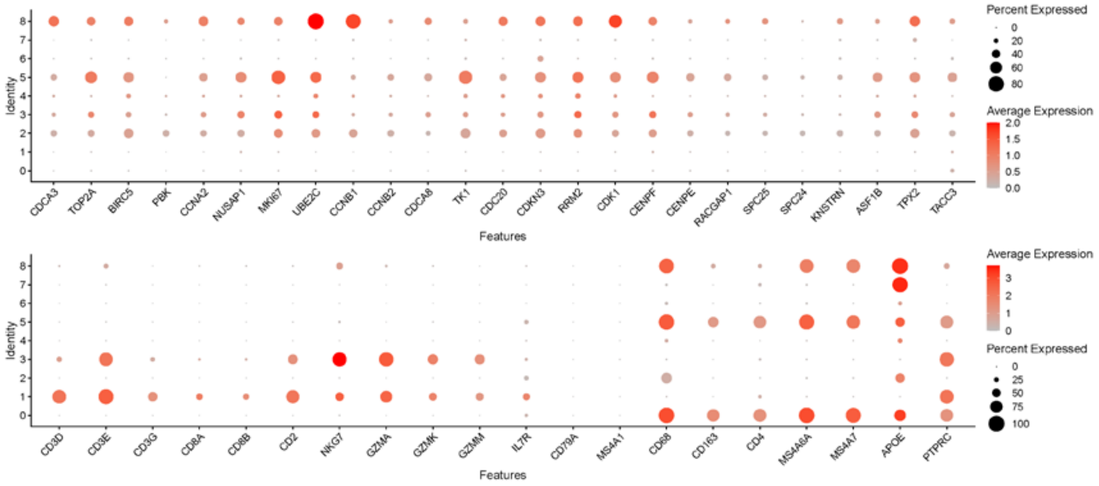


**Fig S4. Nine groups identified using cell-specific markers**

Nine cell groups (GEO: GSM5354533), numbered from 0 to 8, correspond to those in **Figure 1** on a one-to-one basis. Group 3 is predominantly composed of DNtCLCs, while Group 1 includes DN and CD8+ T cells. Group 0 is predominantly composed of OGCs, while Group 5 and 8 are predominantly composed of oCMCs.

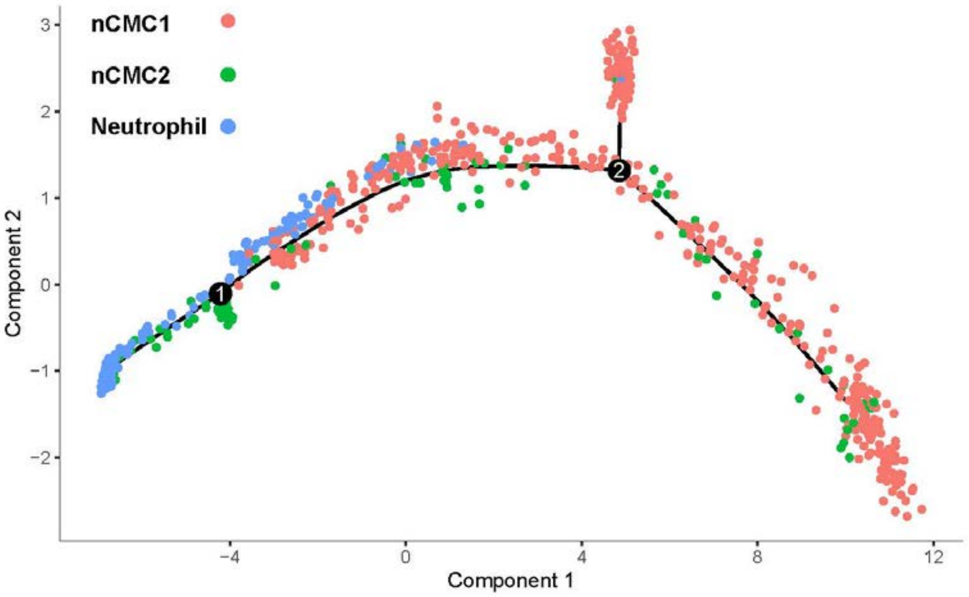


**Fig S5. Nine groups identified using cell-specific markers**

Subset 1 from 49,973 cells (GEO: GSE180286) contained 946 cells from the nCMC1, nCMC2 and neutrophil groups. A linear differentiation path was reconstructed using the top 497 highly variable genes.

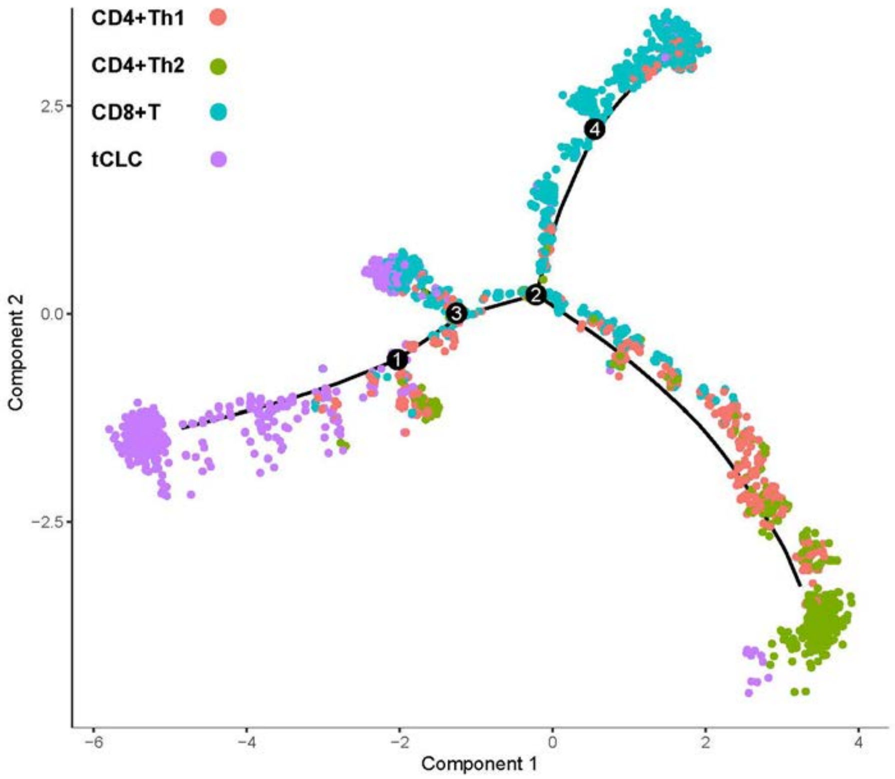


**Fig S6. Nine groups identified using cell-specific markers**

Subset 2 from 49,973 cells (GEO: GSE180286) contained 2,698 cells from the tCLC, CD8+TC, CD4+Th1, and CD4+Th2 groups. A linear differentiation path was reconstructed using the top 2,000 highly variable genes.

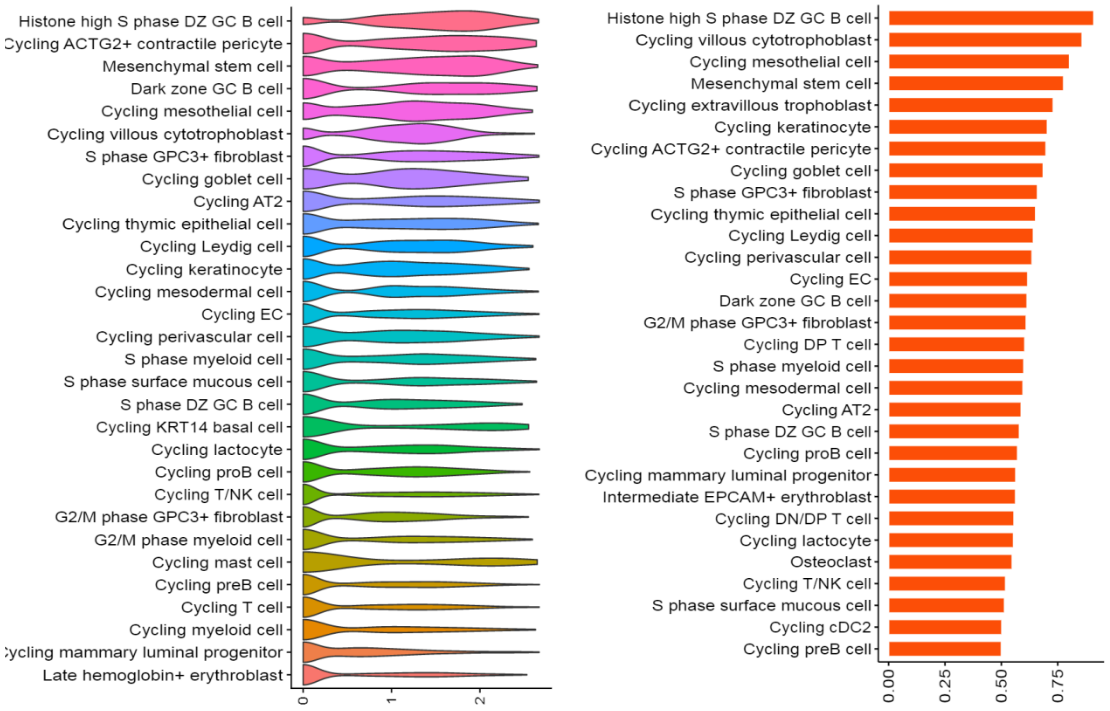


**Fig S7. TK1 gene expression levels in various types of cycling cells.**

The left panel shows the top 30 cell types by mean expression, and the right shows the top 30 by expression percentage. Statistics were computed with data from the DISCO database (https://www.immunesinglecell.org/). Cell-type identities were assigned by DISCO. 26 marker genes are specifically expressed in cycling cells. Here only the TK1 results are shown.

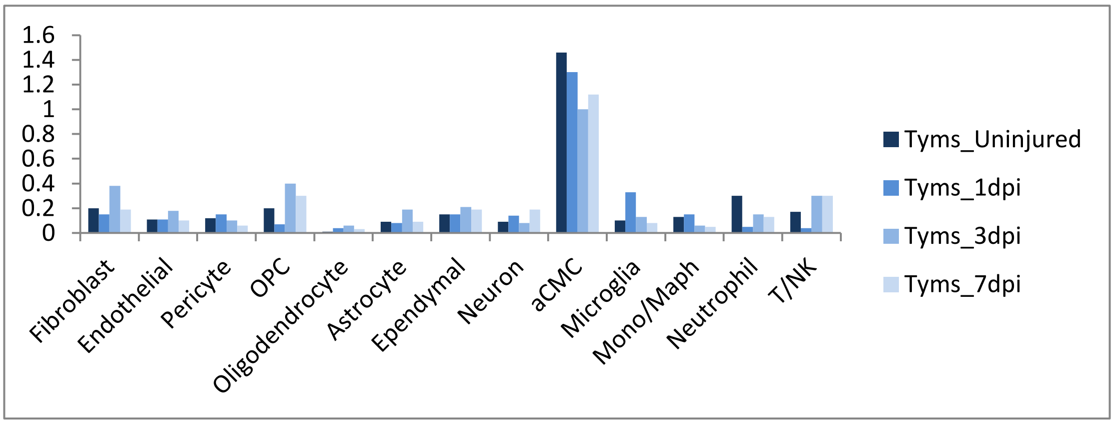


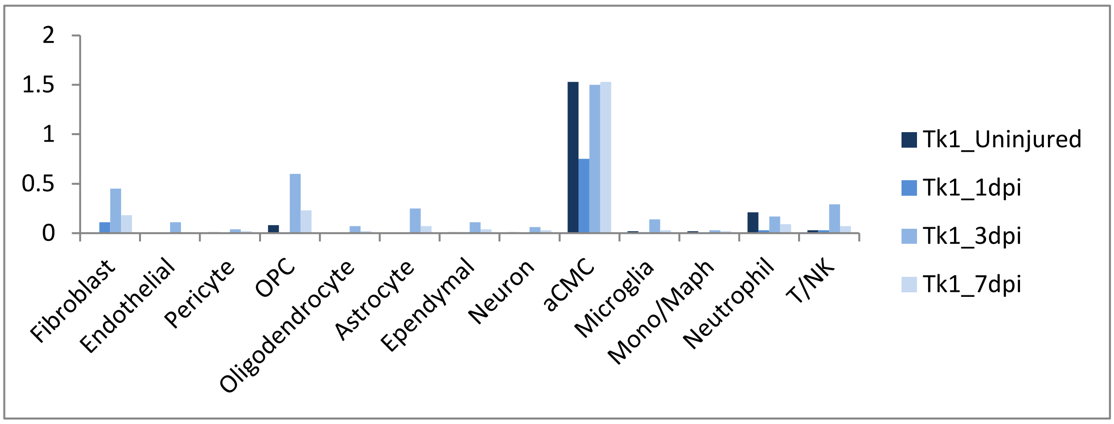

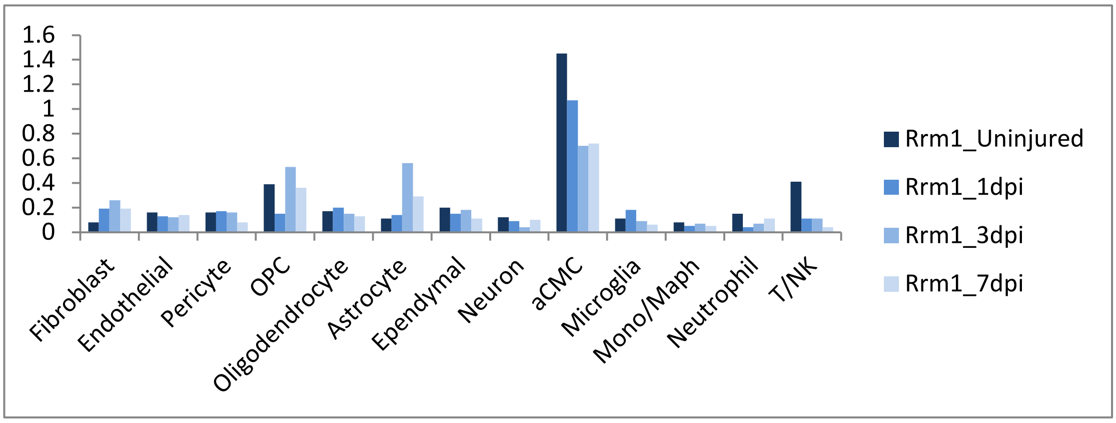
1.6
1.4
1.2
1
0.8
0.6
0.4
0.2
0
Fibroblast
Endothelial
Pericyte
OPC
Oligodendrocyte
Astrocyte
Ependymal
Neuron
aCMC
Microglia
Mono/Maph
Neutrophil
T/NK
Rrm1_Uninjured
Rrm1_1dpi
Rrm1_3dpi
Rrm1_7dpi

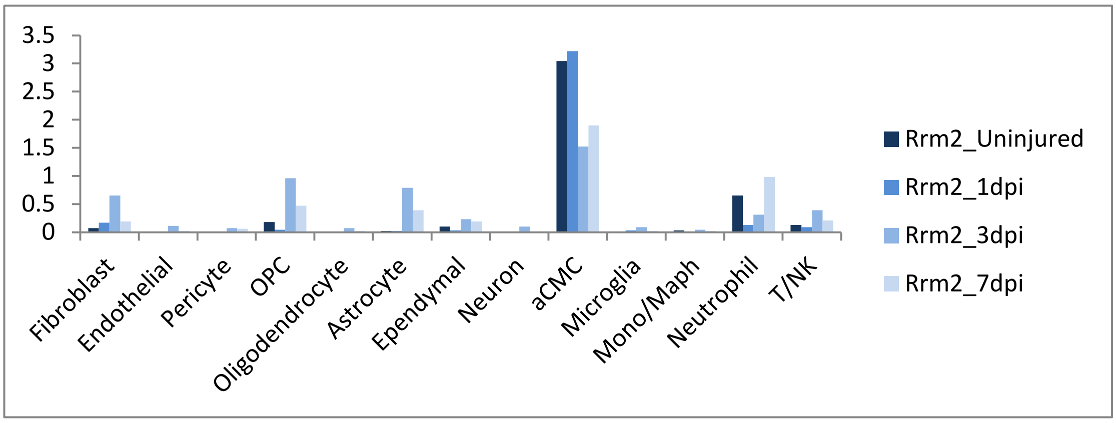
3.5
3
2.5
2
1.5
1
0.5
0
Fibroblast
Endothelial
Pericyte
OPC
Oligodendrocyte
Astrocyte
Ependymal
Neuron
aCMC
Microglia
Mono/Maph
Neutrophil
T/NK
Rrm2_Uninjured
Rrm2_1dpi
Rrm2_3dpi
Rrm2_7dpi

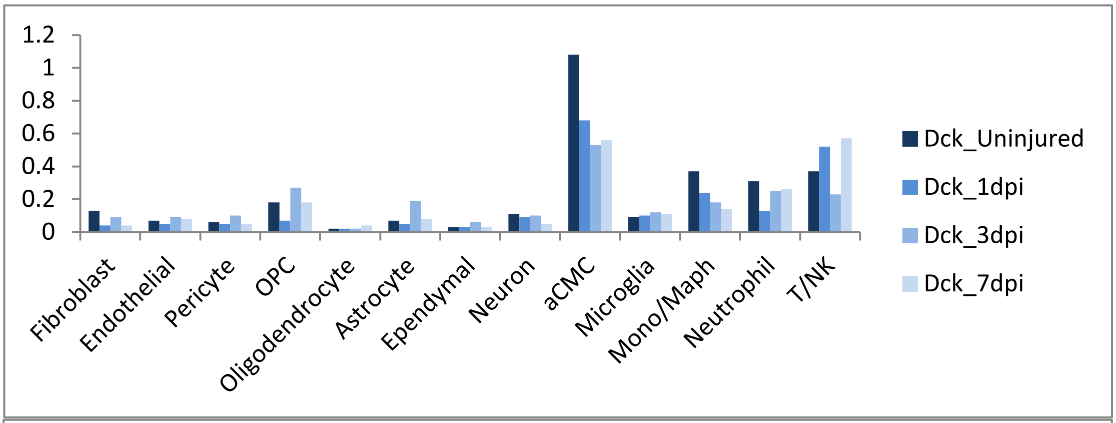
1.2
1
0.8
0.6
0.4
0.2
0
Fibroblast
Endothelial
Pericyte
OPC
Oligodendrocyte
Astrocyte
Ependymal
Neuron
aCMC
Microglia
Mono/Maph
Neutrophil
T/NK
Dck_Uninjured
Dck_1dpi
Dck_3dpi
Dck_7dpi

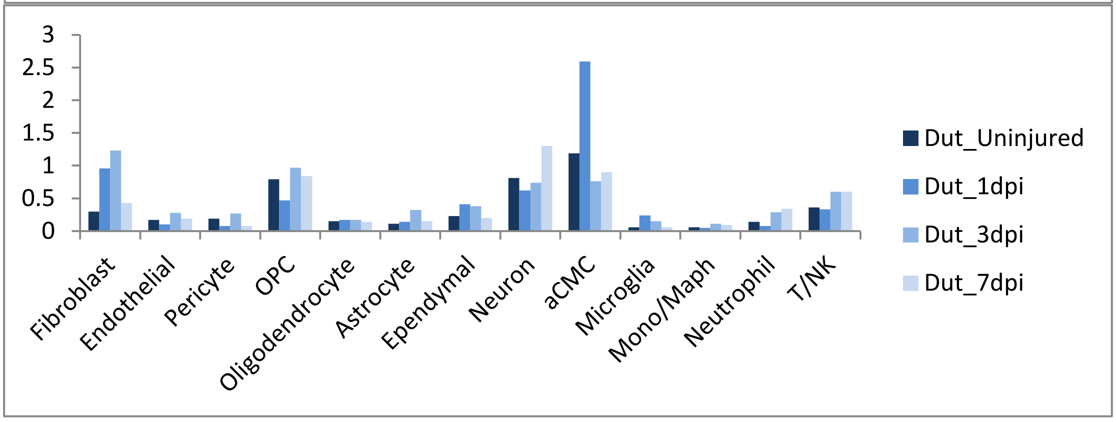
3
2.5
2
1.5
1
0.5
0
Fibroblast
Endothelial
Pericyte
OPC
Oligodendrocyte
Astrocyte
Ependymal
Neuron
aCMC
Microglia
Mono/Maph
Neutrophil
T/NK
Dut_Uninjured
Dut_1dpi
Dut_3dpi
Dut_7dpi

**Fig S8A. Up-regulated expression of six genes in aCMCs**

In our previous study, 66,178 cells from mouse spinal cords were clustered into 13 major groups by re-analysis of a scRNA-seq dataset (GEO: GSE162610). The expression of the key enzymes in the dNTP-synthesis pathways, such as *TYMS*, *TK1*, *RRM1*, *RRM2*, *DCK* and *DUT*, are coordinately up-regulated in aCMCs, compared to other major groups. For each major group, expression data were obtained from uninjured and 1 3 7 day post-injury. Mono/Maph mainly includes monocytes and macrophages, while T/NK includes T lymphocytes, NK cells, etc. OPC: oligodendrocyte precursor cell; aCMC, cycling myeloid cell type a of CMC; Dpi, day post-injury.

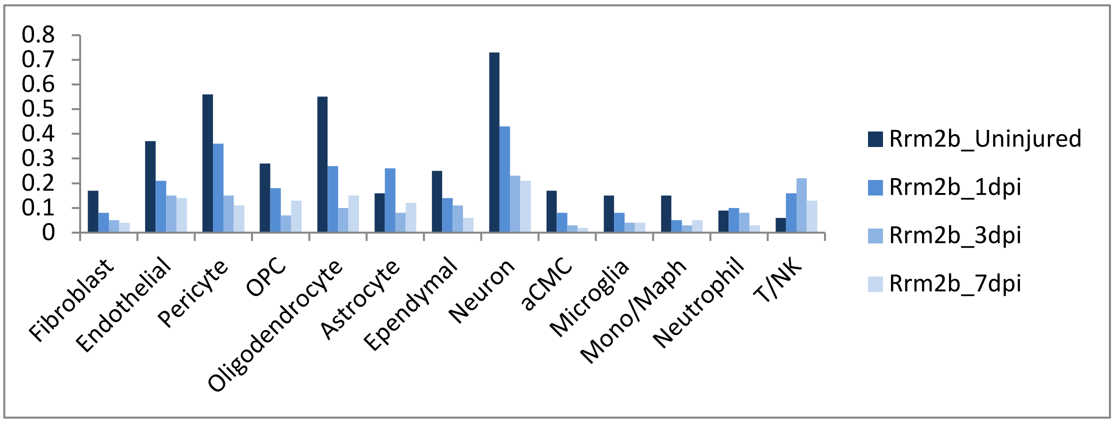


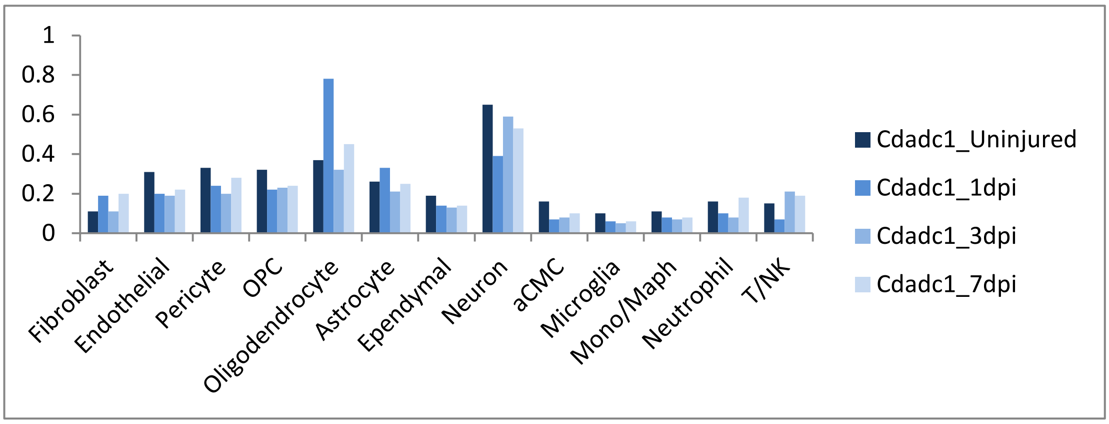


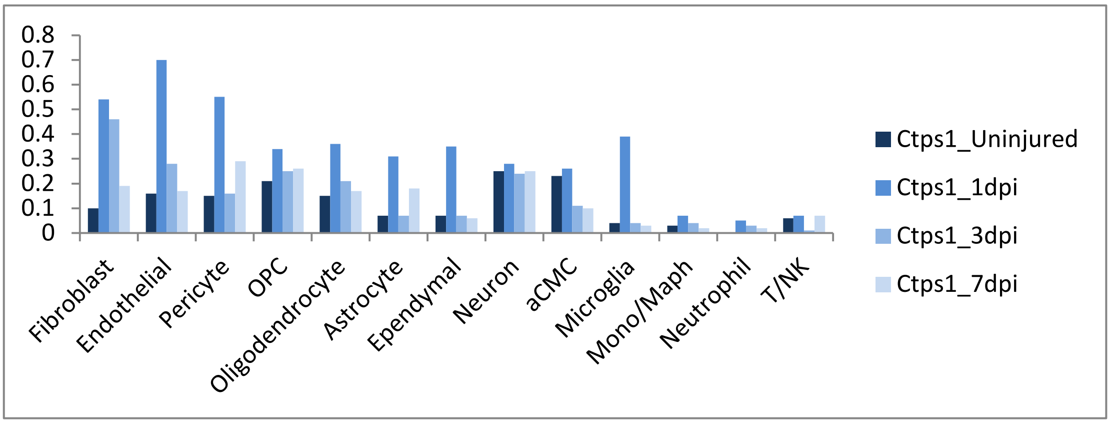

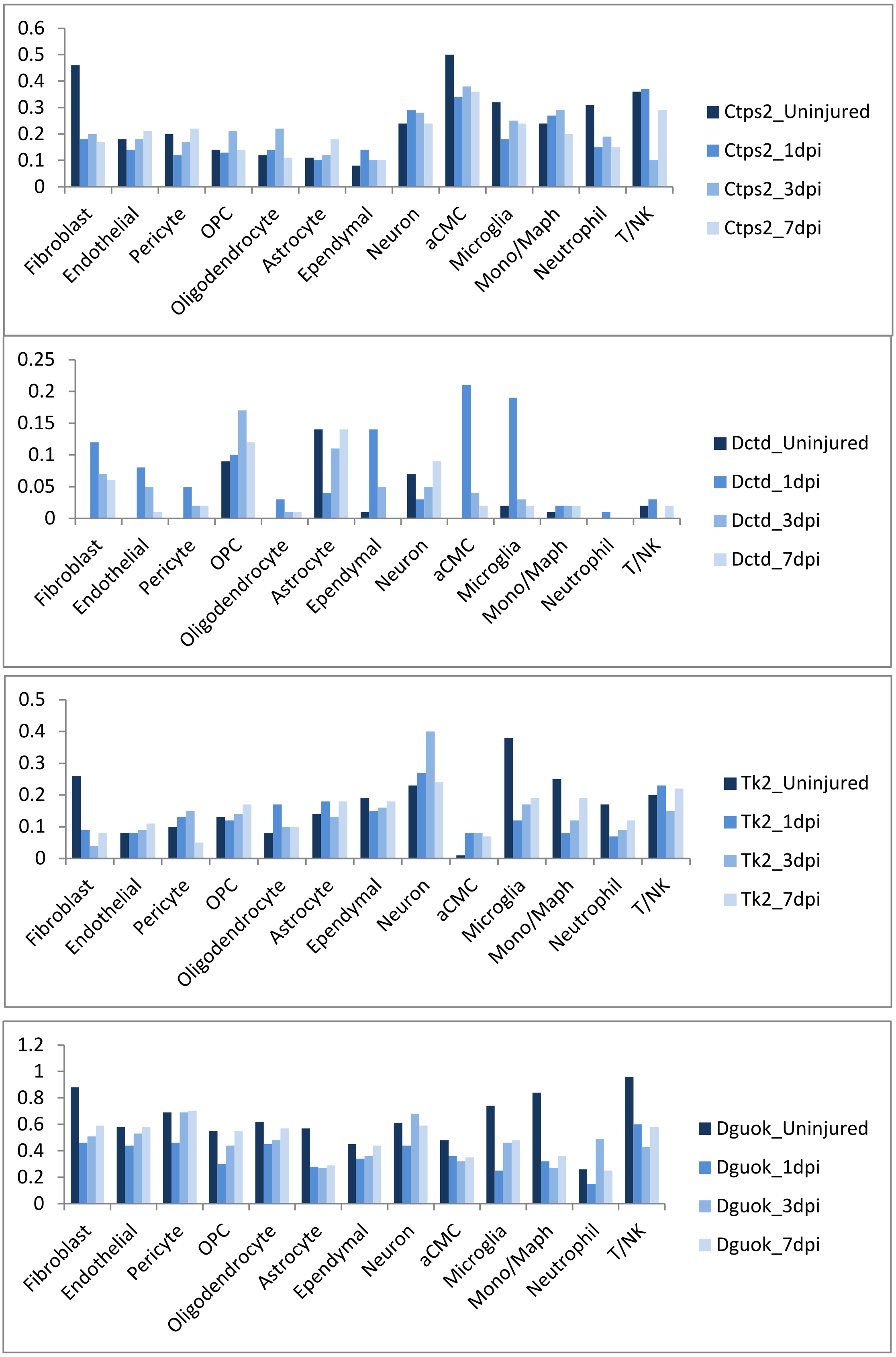


**Fig S8B. Up-regulated expression of seven genes in aCMCs**

The expression of the key enzymes in the dNTP-synthesis pathways, such as *RRM2B*, *CDADC1*, *CTPS1*, *CTPS2*, *DCTD*, *TK2* and *DGUOK*, are not coordinately up-regulated in aCMCs, compared to other major groups. CTPS1/2: CTP synthase 1/2; CMK (*CDADC1*): cytidine/uridine monophosphate kinase 1; RNR: ribonucleotide reductase; DCTD: deoxycytidylate deaminase; DCK: deoxycytidine kinase; DUT: dUTP diphosphatase; TK1/2: thymidine kinase 1/2; TS (*TYMS*): thymidylate synthase; DGUOK: deoxyguanosine kinase. RNR is encoded by *RRM1*, *RRM2* and *RRM2B*.
